# Cell-penetrating peptide-mediated mouse oocyte activation

**DOI:** 10.1101/2025.02.16.638551

**Authors:** Toru Suzuki

**Author notes:** **Correspondence:** Toru Suzuki.

## Abstract

**Introduction:** The cell cycle of ovulated oocytes from various animal species, including mice, is arrested in the second meiotic metaphase until fertilization. The meiotic cell cycle must be initiated to initiate embryonic development. Besides natural fertilization, several other methods have been developed to activate unfertilized oocytes without using sperm. These methods are not only utilized for animal production but have also proven effective in studying the molecular mechanisms that regulate the meiotic cell cycle, oocyte activation, and embryonic development. This study aimed to develop a method to activate mouse oocytes using a cell-penetrating peptide based on the knowledge that the C-terminal domain of the meiotic protein Emi2 can resume the arrested meiotic cell cycle.

**Methods:** This study used female B6D2F1 mice to investigate the effects of a CPP-fused Emi2 peptide on oocyte activation. Second meiotic metaphase oocytes were collected, cultured, and treated with the peptide or strontium chloride. Pronuclear formation, second polar body extrusion, and blastocyst development were assessed, and statistical significance was determined using Fisher’s exact test.

**Results:** The cell-penetrating peptide activated zona-intact oocytes in a manner dependent on specific amino acid residues and peptide concentrations, which are critical components for cell membrane penetration. It has also been confirmed that oocytes activated using this method can develop to the blastocyst stage.

**Discussion:** The introduction of peptides or functional amino acid sequences using CPP or related methods could be an alternative for easily performing functional analyses of the activity of target proteins in oocytes.

## 1 Introduction

In unfertilized oocytes of various species, including mice and frogs, the cell cycle is arrested at the second meiotic metaphase (MII) until fertilization (Sagata, 1996). Therefore, one of the earliest and most critical events in embryogenesis initiation is the resumption of cell cycle arrest in oocytes. Meiotic arrest and resumption are tightly regulated processes, and any disruption in this regulatory mechanism can severely impair the developmental rate of embryos, potentially leading to developmental arrest or infertility in the most severe cases (Hashimoto et al., 1994; Colledge et al., 1994; Backs et al., 2010; Oh et al., 2011; Nozawa et al., 2023).

The molecular mechanisms that regulate MII arrest and meiotic resumption in unfertilized oocytes have been extensively studied in frogs and mice, and a series of molecular events leading to meiotic resumption have been proposed. The transition from metaphase II to anaphase II in oocytes requires the degradation of cyclin, a maturation promoting factor/M-phase promoting factor (MPF) component, with the assistance of an anaphase-promoting complex (APC) with ubiquitin ligase activity, which is inhibited by Emi2, a zinc finger protein (Tung et al., 2005; Shoji et al., 2006), in unfertilized oocytes. Upon fusion of unfertilized oocytes with sperm, the free calcium ion concentration in the oocyte cytoplasm increases, activating CaMKIIγ, a calcium/calmodulindependent kinase (Liu and Maller, 2005). Phosphorylation of Emi2 by activated CaMKIIγ, followed by phosphorylation of the Emi2 degron by Plk1, results in Emi2 degradation and inactivation. The inactivation of Emi2 enhances APC activity, leading to cyclin ubiquitination and degradation and MPF inactivation, ultimately triggering meiotic cell cycle resumption, that is oocyte activation (Liu and Maller, 2005; Madgwick and Jones, 2007).

Besides sperm-mediated oocyte activation, various techniques have been developed for mammalian species to regulate the meiotic cell cycle and artificially stimulate unfertilized oocytes to begin embryogenesis (Nasr-Esfahani et al., 2010). Ethanol (Cuthbertson, 1983), strontium (Whittingham and Siracusa, 1978), and calcium ionophores (Steinhardt and Epel, 1974) are well-known agents used for oocyte activation. Oocyte activation by these agents is associated with molecular events triggered by elevated ooplasmic calcium concentrations that mimic natural fertilization (Fulton and Whittingham, 1978; Fulton and Whittingham, 1978; Cuthbertson et al., 1981). Some oocyte activation methods are employed not only in basic biological research using animal models but also in assisted reproduction technologies to support human reproduction in clinical settings (Tesarik and Sousa, 1995).

Ohe et al. (2010) and Suzuki et al. (2010a) reported that the C-terminal amino acids of Emi2 are essential for inhibiting APC activity in frog and mouse oocytes, respectively. They also demonstrated that the treatment of frog egg extracts with the Emi2 C-terminal peptide (Ohe et al., 2010) or injection of mRNA encoding the Emi2 C-terminal region into unfertilized mouse oocytes (Suzuki et al., 2010a) resumed MII arrest. Based on the observation that the Emi2 C-terminus binds to APC or its component Cdc20 (Ohe et al., 2010; Suzuki et al., 2010a), the Emi2 C-terminal sequence has been proposed to competitively inhibit the binding of endogenous Emi2 to APC in unfertilized oocytes, thereby inducing cell cycle reentry through APC activation and inactivation of M-phase cyclins mediated by endogenous Emi2 inhibition (Ohe et al., 2010; Suzuki et al., 2010a).

Ohe et al. and Suzuki et al. suggested that the Emi2 C-terminal peptide could serve as a novel oocyte activator for mammalian embryo production. However, it remains unclear whether peptides can activate living and intact MII oocytes without inducing undesirable cellular effects that disrupt normal meiotic processes and preimplantation embryo development. To test this hypothesis, it was necessary to introduce the peptide into unfertilized oocytes. Micromanipulation is one of the most effective techniques for introducing molecules into oocytes and embryonic cells (Gordon and Ruddle, 1981; Kimura and Yanagimachi, 1995; Suzuki and Perry, 2018); however, it requires specialized equipment and expertise. Another alternative is the use of cell-penetrating peptides (CPPs). CPPs were first discovered by Frankel and Pabo when studying the HIV viral protein TAT (Frankel and Pabo, 1988). When purified TAT is added to culture media, it is taken up by cultured cells *in vitro*, localized to the nucleus, and activates the viral promoter (Frankel and Pabo, 1988). The CPP sequence of TAT comprises only 13 amino acids (TAT (48-60), Vivès et al., 1997). Since then, several other amino acid sequences of known proteins and artificially designed peptides have been shown to exhibit cell-penetrating activity (Guidotti et al., 2017). Some CPPs contain as few as 13 amino acids and can be chemically synthesized by fusion with target peptides with relative ease.

In this study, the ability of CPP-fused mouse Emi2 C-terminal peptide to induce meiotic resumption in unfertilized mouse oocytes was investigated. These results demonstrated that mouse oocytes with a zona pellucida treated with the CPP-Emi2 C-terminal peptide initiated embryonic development, with some oocytes developing to the blastocyst stage.

## 2 Materials and methods

### 2.1 Animals

All the animal experiments were approved by the Institutional Animal Care and Use Committee of the Institute of Science Tokyo, Japan. Female B6D2F1 mice were obtained from CLEA Japan and maintained under a 12-hour light/dark cycle.

### 2.2 Chemicals

The CPP-fused Emi2 C-terminal peptide contained the last 18 amino acids of mouse Emi2 and eight arginine residues (R8) at its N-terminus. In the AA mutant, the last 2 residues, arginine and leucine, of Emi2 were replaced with two alanine residues. Peptides were obtained from ABclonal Biotechnology K.K. (Massachusetts, US) or GL Biochem (Shanghai) Ltd. (Shanghai, China), dissolved in sterilized water (NAKALAI, Kyoto, Japan) at the stock concentration of 30 mM, and stored at -80 °C until use.

### 2.3 Oocyte collection, *in vitro* culture, and activation

Female B6D2F1 mice, 8–12 weeks of age, were superovulated by sequential injections of 7.5 IU pregnant mare serum gonadotropin (PMSG; Asuka Animal Health, Tokyo, Japan) and 7.5 IU human chorionic gonadotropin (hCG; Asuka Animal Health, Tokyo, Japan) 48 h apart. Then, 15–16 h after hCG injection, the mice were sacrificed, and oviducts containing the cumulus-oocyte complexes were collected. Denuded MII oocytes with degenerated first polar body (Suzuki et al., 2016) were washed and cultured in KSOM medium (ARK Resource, Kumamoto Japan) under 5% CO_2_ at 37 °C after cumulus cell removal using hyaluronidase treatment in M2 medium (Merck, Darmstadt, Germany). For the injection experiments, peptides at the indicated concentrations were injected into MII oocytes in an M2 medium using a piezo-actuated micromanipulator (PMM-150, Primetech, Ibaraki, Japan). Physical incision of the zona pellucida of MII oocytes was performed in an M2 medium using a glass needle attached to a micromanipulator (Narishige, Tokyo, Japan). For peptide treatment, zona-intact and incised MII oocytes were cultured in KSOM containing the indicated concentrations of peptides, with or without 5 μg/mL cytochalasin B (FUJIFILM Wako, Tokyo, Japan), for 2.5 h. MII oocytes were treated with 5 mM strontium chloride (Merck, Darmstadt, Germany) in a calcium-free CZBG medium containing cytochalasin B for 2.5 h to induce diploid parthenogenotes. After treatment with a cytochalasin B-containing medium, peptide- or strontium-treated MII oocytes were further cultured in KSOM with 5 μg/mL cytochalasin B for 4.5 h. The resulting embryos were washed and cultured in KSOM for 5 d. Pronuclear formation and second polar body extrusion were observed at 6–7 h after oocyte activation. Blastocyst development was assessed 5 d after oocyte activation.

### 2.4 Statistics

Fisher’s exact test was performed, and p-values less than 0.05 were considered significantly different. All experiments were conducted over more than two separate days.

## 3 Results

### 3.1 Design and activity of Emi2 C-terminal peptide

The mouse Emi2 protein contains a highly conserved RL tail domain at its C-terminus in several species, including Xenopus laevis (Suzuki et al., 2010a; Ohe et al., 2010). It has been reported that a peptide consisting of 14 amino acids from the RL tail of the Xenopus Emi2/Erp1 protein can resume the meiotic cell cycle in Xenopus egg extract (Ohe et al., 2010). Additionally, the mRNA encoding residues 551–641 of the mouse Emi2 protein activates mouse MII oocytes (Suzuki et al., 2010a). Based on this information, I suggest that the conserved RL tail of mouse Emi2 can activate mouse MII oocytes, thereby promoting pre-implantation embryo formation.

To test this possibility, a peptide sequence containing 18 residues from the C-terminal end of the mouse Emi2 protein, along with a cell-penetrating peptide (CPP) and 8 N-terminal arginines (R8; Mitchell et al., 2000; Futaki et al., 2001) was designed and synthesized as a putative cell-penetrating oocyte activation peptide (R8-RL peptide). Because most C-terminal arginine and leucine residues of the Emi2 protein are essential for the function of the RL tail (Ohe et al., 2010; Suzuki et al., 2010a), a mutant peptide in which these two residues (RL) were substituted with two alanine residues (AA) was synthesized as a negative control (R8-AA peptide).

First, these peptides were directly injected into MII oocytes by micromanipulation to analyze their effects on meiotic arrest (Supplementary Table S1). At a concentration of 30 mM, the R8-RL peptide activated 92% of surviving oocytes (n = 25), as evidenced by pronuclear formation, which was significantly different (p = 2.3 × 10^−11^) from the R8-AA peptide (5.7%, n = 35). At lower peptide concentrations, the R8-RL peptide did not show significant differences in oocyte activation compared to the control. These results suggested that the R8-RL peptide has concentration- and RL residue-dependent activities that induce mouse oocyte activation.

### 3.2 Zona intact oocytes are activated with RL peptide

CPP has been reported to move across the cell membrane into the cell cytoplasm (Frankel and Pabo, 1988; Vivès et al., 1997); however, whether the peptide designed in this study crosses the zona pellucida, an extracellular matrix, to access the oocytes is unknown. To investigate this, MII oocytes with intact or severed zona pellucida were cultured in a culture medium containing 3 mM RL peptide for 6 h (Table 1). The R8-RL peptide at the concentration of 3 mM activated not only the surviving MII oocytes with severed zona pellucida with a micro glass needle (100%, n=9) but also those with intact zona pellucida (100%, n=7) with a significant difference from the R8-AA peptide (p=8.0×10^-8^ for zona-intact oocytes and 8.4×10^-6^ for zona-cut oocytes, RL vs AA). Some MII oocytes died after treatment with these peptides in both the intact and zona-cut groups, indicating their cytotoxicity. These results show that the CPP-conjugated Emi2 RL peptide composed of 26 amino acids used in this study (R8-RL) can activate MII oocytes, regardless of the presence or absence of an intact zona pellucida.

**Table 1.** Oocyte activation after the treatment with cell-penetrating peptides, with or without zona pellucida.

| Peptide concentration (mM) | 3 |  |  |  | 0.3 |  |  |  |
| --- | --- | --- | --- | --- | --- | --- | --- | --- |
| Zona | Intact |  | Cut |  | Intact |  | Cut |  |
| Emi2 C-terminal | RL | AA | RL | AA | RL | AA | RL | AA |
| Total number of oocytes | 37 | 37 | 35 | 36 | 37 | 37 | 35 | 37 |
| Number of dead oocytes | 28 | 9 | 28 | 10 | 14 | 0 | 12 | 0 |
| Number of surviving oocytes | 9 | 28 | 7 | 26 | 23 | 37 | 23 | 37 |
| Number of activated oocytes | 9* | 1 | 7* | 2 | 3 | 0 | 1 | 0 |
| Number of MII oocytes | 0 | 27 | 0 | 24 | 20 | 37 | 22 | 37 |

\*p<0.05, RL vs AA.
Fisher's exact test was performed on the activated oocytes within the surviving oocytes.

### 3.3 *In vitro* development of MII oocytes activated with RL peptide

The *in vitro* developmental capacity of oocytes activated by R8-RL peptide was examined. All MII oocytes treated with the R8-RL peptide at any concentration tested (3 to 0.5 mM) survived after 1 h of treatment, but up to 31.4% were dead after 2.5 h (Supplementary Table S2). All of the concentrations from 3 to 0.5 mM tested in this study activated a part of surviving MII oocytes (39.1 to 79.4% of surviving oocytes, n=34 to 68). Of the activated oocytes, 23.5–42.6% had a second polar body, and 22.2–55.6% were developed to the blastocyst stage after 5 d of *in vitro* culture (Table 2).

**Table 2.** Oocyte activation and *in vitro* development after the peptide treatment.

| Peptide concentration (mM) | 3 | 2 | 1 | 0.75 | 0.5 | 0 |
| --- | --- | --- | --- | --- | --- | --- |
| Total number of oocytes | 53 | 49 | 98 | 48 | 50 | 57 |
| Number of dead oocytes (%)<sup>*1</sup> | 19 (35.8) | 10 (20.4) | 30 (30.6) | 8 (16.7) | 4 (8.0) | 0 (0.0) |
| Number of surviving oocytes | 34 | 39 | 68 | 40 | 46 | 57 |
| Number of pronuclear formation (%)<sup>*2</sup> | 27 (79.4) | 27 (69.2) | 34 (50.0) | 26 (65.0) | 18 (39.1) | 0 (0.0) |
| Number of pronuclear formation with 2<sup>nd</sup> polar body (%) | 8 (23.5) | 13 (33.3) | 29 (42.6) | 17 (42.5) | 15 (32.6) | 0 (0.0) |
| Number of MII oocytes | 7 | 12 | 34 | 14 | 28 | 57 |
| Number of Blastocyst (%)<sup>*3</sup> | 6 (22.2) | 15 (55.6) | 11 (32.4) | 11 (42.3) | 9 (50.0) | 0 (0.0) |

<sup>\*1</sup>% of Total.
<sup>\*2</sup>% of surviving oocytes.
<sup>\*3</sup>% of pronuclear embryos.

Activated oocytes from the preceding experiment were assumed to be a mixture of haploid and diploid parthenogenotes based on the observation that they contained pronuclei with or without a second polar body. To analyze the developmental potential of diploid embryos, MII oocytes were treated with R8-RL peptide in the presence of an actin polymerization inhibitor, cytochalasin B, to block polar body extrusion and produce diploid embryos. It was initially confirmed (Supplementary Table S3) that 2 and 3 mM of RL peptides without R8 residues did not activate surviving zona-intact MII oocytes (0% activation, n = 50 for each concentration). After treatment with 2 to 0.5 mM R8-RL peptide, 47.9–75.0% of surviving oocytes (n = 27–48) formed pronuclei but did not exhibit a second polar body, suggesting the production of diploid parthenotes (Table 3). Of the activated oocytes, 7.1– 87.0% developed to the blastocyst stage. A total of 92% of the oocytes treated with both strontium and cytochalasin B (n = 50) were activated and 100% of them developed to the blastocyst stage. These results indicate that mouse pre-implantation embryos produced by R8-RL have the potential to develop into the blastocyst stage.

**Table 3.** Oocyte activation and *in vitro* development after the peptide treatment in the presence of CB.

| Peptide concentration (mM) or Sr | Sr | 2 | 1 | 0.5 | 0 |
| --- | --- | --- | --- | --- | --- |
| Total number of oocytes | 50 | 50 | 50 | 50 | 30 |
| Number of dead oocytes (%)<sup>*1</sup> | 4 (8.0) | 23 (46.9) | 18 (36.0) | 2 (4.9) | 0 (0.0) |
| Number of surviving oocytes | 46 | 27 | 32 | 48 | 30 |
| Number of MII oocytes | 0 | 13 | 8 | 25 | 30 |
| Number of pronuclear formation (%)<sup>*2</sup> | 46 (100.0) | 14 (51.9) | 24 (75.0) | 23 (47.9) | 0 (0.0) |
| Number of Blastocyst (%)<sup>*3</sup> | 46 (100.0) | 1 (7.1) | 10 (41.6) | 20 (87.0) | 0 (0.0) |

<sup>\*1</sup>% of Total.
<sup>\*2</sup>% of surviving oocytes.
<sup>\*3</sup>% of pronuclear embryos.

## 4 Discussion

This study demonstrated that the CPP (R8)-conjugated mouse Emi2 RL domain peptide can activate zona-intact mouse MII oocytes in a CPP- and RL residue-dependent manner, leading to the formation of blastocyst-stage embryos.

CPPs have been identified in several cell-penetrating proteins and have been investigated to introduce various molecules that control cellular functions in living cells. This technique has already been applied in studies using unfertilized mammalian oocytes and preimplantation embryos (Kwon et al., 2009; Kwon et al., 2013; Klinsky et al., 2023), but reports vary as to whether sufficient transduction efficiency can be achieved. It has been reported that CPP-conjugated LDP12–EGFP–LC3 is not incorporated into mouse MII oocytes enough to detect fluorescent signals in ooplasm (Kwon et al., 2013). Another report showed that a CPP-conjugated protein (a permeable tetanus toxin light chain, CPP-TeTx) can move into mouse oocytes regardless of the presence of the zona pellucida, leading to the expected inhibition of cortical granule exocytosis during fertilization (Klinsky et al., 2023). The number of amino acids constituting the cargo protein introduced into the oocytes, the concentration of CPP proteins, and the CPP sequences may have a significant effect on the efficiency of intracellular transduction by CPP. Considering that concentrations of 0.5 mM or higher were required for the Emi2 RL peptide (18 amino acids) to activate unfertilized mouse oocytes, and, likely, the introduction of a sufficient number of molecules by CPP in the cell depends on the nature of the cargo being introduced. Some proteins or peptides conjugated to CPP, such as TeTx, exhibit sufficient activity at lower concentrations (4 μM) to regulate cellular functions. However, other compounds display reduced activity, necessitating higher concentrations to effectively regulate target molecules, which can result in increased cytotoxicity. It is important to find a way to control the cytotoxicity of CPP-cargo conjugates or increase their activity in cells (Reissmann, 2014; Kim et al., 2021), which is an obstacle to overcome for their widespread use in cell and/or fertilization research. Since it was shown in this study that peptide concentration and treatment time are critical parameters for higher survival and development rates of treated oocytes, their adjustment is necessary to establish better experimental conditions. Some peptides, including Emi2 RL, would require higher concentrations to act on oocytes to regulate cellular processes. Additionally, it may be beneficial to use other CPP sequences with lower toxicity, substances that promote the intracellular delivery of peptides, and reagents that facilitate the repair of damaged cell membranes during CPP penetration.

The results in this study indicate that the Emi2 RL domain peptide triggers sufficient activity to activate MII oocytes and produce blastocyst-stage embryos. Previous reports (Suzuki et al., 2010a; Ohe et al., 2010) suggest that the Emi2 RL peptide introduced into unfertilized mouse oocytes functions as a negative endogenous Emi2 regulator, which is sufficient to produce embryos. It has previously been shown that a zinc ion chelator, TPEN, can also activate mouse MII oocytes (Suzuki et al., 2010b; Kim et al., 2010). The Emi2 protein has a zinc finger domain that is thought to be essential for Emi2 as an APC/C inactivator and has been proposed to remove zinc from MII oocytes, leading to meiotic resumption through Emi2 inactivation and APC/C activation. The proposed mechanism of meiotic resumption is similar to treatment with the Emi2 RL peptide and TPEN in terms of Emi2 function inactivation. The developmental potential from the 1-cell to blastocyst stage of parthenogenetic diploid embryos produced by the Emi2 RL peptide in this study was comparable to that produced by TPEN in a previous study (Suzuki et al., 2010b). The direct functional regulation of target factors downstream of calcium signaling by RL peptides or TPEN allows for a more detailed analysis of the function of meiotic cell cycle regulatory molecules and fertilization-associated events. For example, the finding that TPEN can induce oocyte activation reveals the essential role of zinc in MII arrest. Furthermore, it has been suggested that the only developmentally essential role of calcium oscillation during fertilization is to induce meiotic resumption, since TPEN-mediated oocyte activation is not associated with an ooplasmic calcium pulse but can be used to produce fertile mice. The activity of the Emi2 RL peptide demonstrated in this study supports the idea that the endogenous mouse Emi2 RL domain, which comprises fewer than 18 amino acids, is involved in the maintenance of MII arrest.

The use of chemical inhibitors is a powerful way to control the function of molecules in cells and reveal their intracellular functions. However, there are many proteins, including Emi2, for which no inhibitors have been reported and the low substrate specificity of these chemical inhibitors can be problematic. Molecular control technologies based on the principle of protein-protein interactions, such as the peptides used in this study, can be relatively easily designed from naturally occurring amino acid sequences based on an understanding of the mechanisms of protein behavior and are expected to have higher substrate and interaction specificity. In cells such as unfertilized oocytes or embryonic cells at the earliest developmental stages, where gene expression has not yet occurred (Moore et al., 1974; Bouniol-Baly et al., 1999; Minami et al., 2007), and some essential proteins have already accumulated in sufficient quantities (Minami et al., 2007) to support the earliest developmental events, genetic depletion, and RNA degradation-inducing techniques often exhibit reduced or no efficiency when attempting to inhibit the function of targeted proteins. The introduction of peptides or functional amino acid sequences using CPP or related methods could be an alternative for easily performing functional analyses of the activity of target proteins in oocytes. However, as demonstrated in this study, CPP-conjugated peptides exhibit cytotoxicity and may interfere with meiotic processes, such as the polar body extrusion of oocytes. Mitigating or eliminating these detrimental effects of CPPs on living oocytes remains a significant challenge.

## 6 Ethics statement

All the animal experiments were approved by the Institutional Animal Care and Use Committee of the Institute of Science Tokyo, Japan.

## 7 Author Contributions

TS: Conceptualization, Formal analysis, funding acquisition, Investigation, Methodology, Writing the original draft, writing the review, and editing.

