## Supplemental Table 1-3 for "Cell-penetrating peptide-mediated mouse oocyte activation"

### *Supplementary Material*

#### 1 Supplementary Tables

Supplementary Table S1. Oocyte activation after peptide injection into mouse unfertilized oocytes.

| Peptide concentration (mM) | 30 | 3 | 0.3 |
|---|---|---|---|
| AA | 2/35 | 1/30 | 2/37 |
| RL | 23/25* | 2/31 | 2/38 |

*$p<0.05$, RL vs AA.

Supplementary Table S2. Effect of treatment duration for oocyte survival.

| Peptide concentration (mM) | 3 | 2 | 1 | 0.75 | 0.5 | 0 |
|---|---|---|---|---|---|---|
| Total number of oocytes | 55 | 55 | 105 | 50 | 50 | 57 |
| Number of dead oocytes at 1 h | 0 | 0 | 0 | 0 | 0 | 0 |
| Number of dead oocytes at 2.5 h (%) | 16 (29.1) | 12 (21.8) | 33 (31.4) | 10 (20.0) | 2 (4.0) | 0 (0.0) |

Supplementary Table S3. Effect of CPP conjugation on oocyte activation.

| Peptide concentration (mM) | 2 | 2 | 3 |
|---|---|---|---|
| CPP conjugation | + | - | - |
| Total number of oocytes | 50 | 50 | 50 |
| Number of dead oocytes (%)[*1] | 19 (38.0) | 0 (0.0) | 0 (0.0) |
| Number of surviving oocytes | 31 | 50 | 50 |
| Number of MII oocytes | 21 | 50 | 50 |
| Number of pronuclear formation (%)[*2] | 10 (32.3) | 0 (0.0) | 0 (0.0) |

[*1]% of Total.

[*2]% of surviving oocytes.
